# Combination antibiotic PMMA composites provide sustained broad-spectrum antimicrobial activity and allow for post-implantation refilling

**DOI:** 10.1101/832402

**Authors:** Erika L. Cyphert, Chao-yi Lu, Dylan W. Marques, Greg D. Learn, Horst A. von Recum

**Affiliations:** Department of Biomedical Engineering, Case Western Reserve University, 10900 Euclid Avenue, Cleveland, OH 44106

**Keywords:** Broad-spectrum, Poly(methyl methacrylate), Polymer, Cyclodextrin, Biomaterial, Infection

## Abstract

Antibiotics are commonly added to poly(methyl methacrylate) (PMMA) by surgeons to locally treat infections such as in bone cement for joint replacement surgeries, but also as implantable antimicrobial “beads”. However, this strategy is of limited value in high risk patients where infections can be recurrent or chronic and otherwise hard to treat. Also when only one drug is incorporated and applied toward poly-microbial infections (multiple bacterial species), there is a high risk that bacteria can develop antibiotic resistance. To combat these limitations, we developed a combination-antibiotic PMMA composite system comprised of rifampicin-filled β-cyclodextrin (β-CD) microparticles added into PMMA filled with a second drug. Different formulations were evaluated through Zone-of-Inhibition, drug activity, antibiotic release and re-filling, as well as mechanical studies. Our combination-antibiotic PMMA composite system achieved up to an eight-fold increase in duration of antimicrobial activity in comparison to clinically used antibiotic-filled PMMA. Inclusion of CD microparticles also allowed for refilling of additional antibiotics after simulated implantation, resulting in additional windows of therapeutic efficacy. Mechanical testing showed that our tested formulations did have a small, but significant decrease in mechanical properties when compared to unmodified controls. While further studies are needed to determine whether the tested formulations are still suitable for load-bearing applications (e.g. bone cement), our composites are certainly amenable for a variety of non-load bearing applications (e.g. antimicrobial “beads” and temporary spacer in two-stage arthroscopic revisions).

## INTRODUCTION

Infections following orthopedic procedures such as total hip and knee arthroplasties have a relatively low occurrence, ranging from 0.3 – 1.28% ^1–3^. Nevertheless, patients that acquire infections often must undergo intensive systemic antibiotic regimens and unpalatable revision surgeries that place a substantial burden on them, both physically and financially ^1, 4–5^. Additionally, revision procedures increase the patient’s risk for further infection, morbidity, and mortality ^1,6^.

The aggressiveness of periprosthetic infections can be exacerbated if the infection is comprised of both gram-positive and gram-negative pathogens. While gram-positive *Staphylococci* species, specifically *Staphylococcus aureus* (*S. aureus*) and *Staphylococcus epidermidis* (*S. epidermidis*) account for the majority of orthopedic infections, up to 15% of infections are caused by gram-negative species ^7^. *Escherichia coli* (*E. coli*) accounts for upwards of 20-30% of gram-negative orthopedic infections ^7^.

In these complex cases, monotherapy antibiotic treatment may not provide broad-spectrum antimicrobial coverage or bacteria may become resistant towards the antibiotic over an extended period of time. In an effort to mitigate the risk of developing drug resistance, antibiotics are often administered in combination ^8–9^. The decreased risk of drug resistance with combination therapies is attributed to the different modes of action of the antibiotics against bacteria, and the increased difficulty for bacterial populations to acquire simultaneous resistance to each unique compound ^9^. For example, one clinical study showed that 42% of patients treated with monotherapy for bacterial infection developed antibiotic-resistant bacteria, while only 17% of patients treated with a combination of two antibiotics developed resistant bacteria ^10^. In addition to decreasing the risk for drug resistance, combination therapies have also been found to be more suitable for treating poly-microbial infections due to their broader antimicrobial coverage ^10–11^. In the clinical study previously mentioned, *Pseudomonas aeruginosa* (*P. aeruginosa*), was the most common gram-negative bacteria species detected in that study; and of the patients studied, *P. aeruginosa*, was found in approximately 19% of patients receiving monotherapy versus only 4% of those receiving combination therapy ^10^. Gribble *et al.* also found that monotherapy had inferior antibacterial activity against *S. aureus* and *S. epidermidis* which resulted in the need for an additional anti-staphylococcal treatment therapy in 38% of patients treated with monotherapy versus only in 29% of those treated with combination therapy ^10^.

Furthermore, combination antibiotic therapy can often result in a synergistic antibacterial effect as a result of complex interactions between the drugs and bacteria. For example, one drug may increase uptake of the other drug by enhancing permeability of the bacterial cell wall. This may explain the synergism observed when aminoglycoside and beta-lactam antibiotics are used in combination ^12^. Another potential mechanism for synergism to occur is to use drugs that target specific metabolic pathways leading to secondary effects that can further enhance the potency of drugs ^12^. This can be observed when trimethoprim and sulfonamides are used against *E. coli* mutants ^12^.

In order to locally deliver antibiotics in orthopedic procedures, such as knee or hip replacements, an antibiotic is often directly added to poly(methyl methacrylate) (PMMA) bone cement during the mixing process ^13^. PMMA is used in a variety of orthopedic and non-orthopedic applications including as a temporary spacer in a two-part revision procedure ^14^, as an anchor between the patient’s bone and the metallic implant in arthroplasties ^15^, and as antibiotic-filled beads ^16–17^. In these applications, typically only one antibiotic, such as gentamicin or tobramycin, is used in an effort to provide antimicrobial coverage ^18^. The primary disadvantages to this strategy are a) these antibiotics may not provide adequate broad-spectrum coverage, b) using only one drug increases risk of developing drug resistance, and c) that the window of antimicrobial activity in these mixtures is limited by the initial amount of antibiotic present within the PMMA, as it is not feasible to replenish the drug reservoir once depleted following implantation ^17,19–20^. Specifically, tobramycin has been found to be less effective in treating *Enterococcus* infections ^19^, and the routine use of gentamicin in PMMA has led to an increased risk of developing bacterial resistance ^20^. Furthermore, there is often a burst release of the antibiotic from the PMMA cement, which results in a suboptimal delivery profile for the treatment of prolonged infections ^13^. While others have developed a variety of strategies such as using liposomes ^21^, adding porogens ^22^, or using calcium phosphate spheres ^23^ to overcome the poor release kinetics of antibiotics from PMMA, there are not any existing delivery systems capable of providing broad-spectrum, long-term, and refillable therapy.

To create a formulation of PMMA bone cement that can effectively treat a range of bacterial infections and that has the ability to be refilled with antibiotics after implantation, we evaluated a composite system comprised of PMMA, combinations of antibiotics, and insoluble β-cyclodextrin microparticles (β-CD). The particles add multiple features including prolonged release and refilling which we have previously shown, as well as the capability of tuning the release rate of one drug separately from the other. Figure 1 depicts a schematic of the composition of the different PMMA composites containing both different drug combinations and differing amounts of β-CD microparticles.

**Figure 1:**
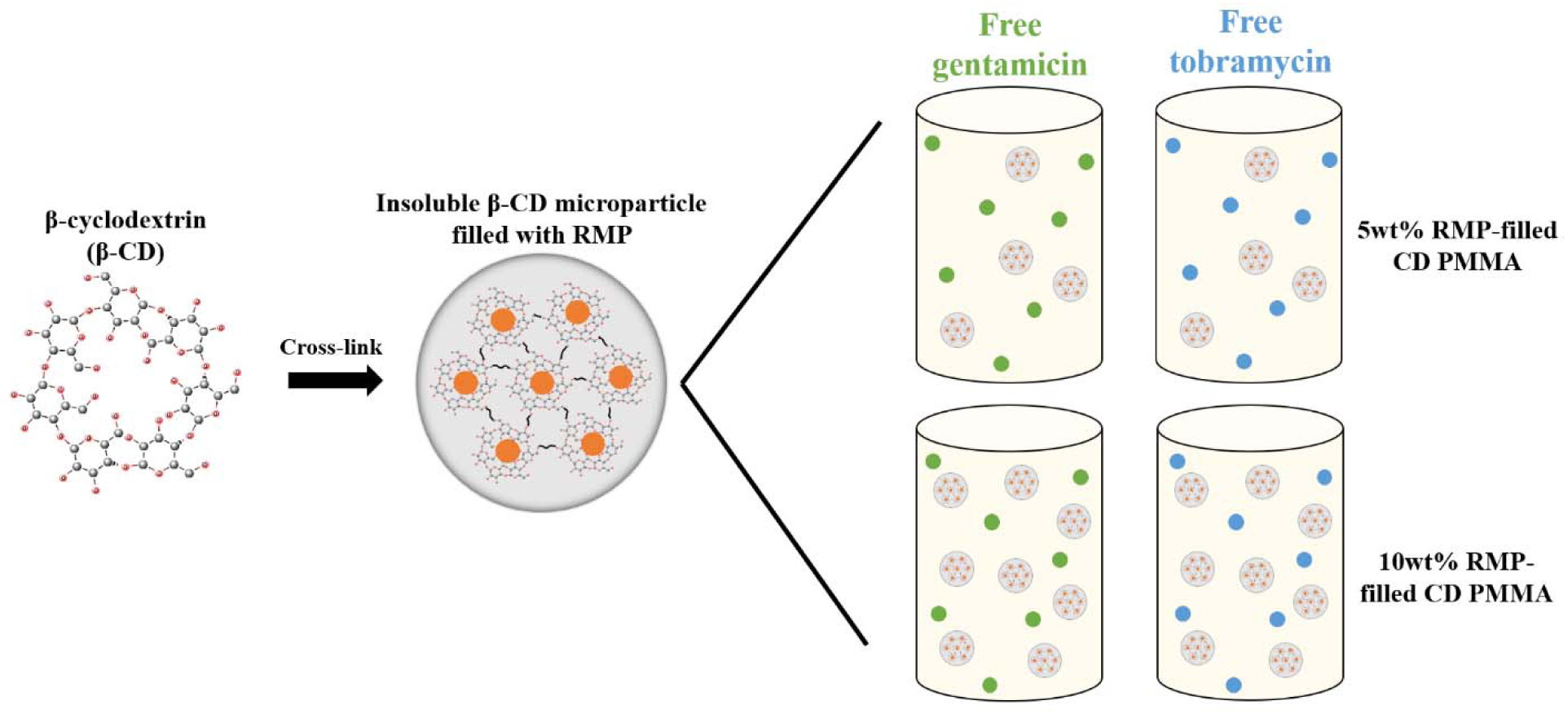
Schematic depicting drug combination composite formulations of PMMA bone cement. Pre-polymerized β-cyclodextrin (β-CD) was cross-linked into insoluble microparticles that were filled with rifampicin (RMP). Different amounts of drug filled β-CD microparticle (either 5 or 10wt%) were incorporated into PMMA during polymerization along with either gentamicin or tobramycin (non-encapsulated in β-CD).

Cyclodextrin is a cyclic oligosaccharide with a hydrophilic exterior and relatively hydrophobic interior. The hydrophobic core allows for an affinity-based interaction with the drug that helps to regulate the release of the drug from the CD cavity ^24^. When additives, such as antibiotics or β-CD microparticles, are used in the PMMA cement it is important to consider how they may affect the mechanical properties of the material. Our previous study has shown that addition of antibiotic-loaded insoluble β-CD microparticles into PMMA cement may mitigate the negative effects on the mechanical properties observed when antibiotic is directly added into PMMA ^17^. Therefore, for this study, we were interested in analyzing the impact of adding the two antibiotics and β-CD microparticles on the mechanical properties of PMMA bone cement. Additionally, in our past study, extended antimicrobial activity was observed after refilling of antibiotics in PMMA with CD microparticles and not in those without microparticles which suggested that the presence of β-CD microparticles in PMMA allowed for an extended therapeutic window for treatment chronic or recurrent infections ^17^. We hypothesized that our PMMA bone cement formulations containing combination antibiotics and β-CD microparticles would allow for an extended antimicrobial effect that is effective against a broader range of bacterial species in comparison to clinically used monotherapy drug formulations.

In this study, we have developed several PMMA bone cement composite formulations containing combination antibiotics (i.e. either gentamicin or tobramycin with rifampicin) with different amounts of β-CD microparticles to evaluate their efficacy in treating broad-spectrum bacterial species as well as their suitability for treating long-term or latent infections. Specifically, we have evaluated the duration of antimicrobial activity of different combination-antibiotic PMMA cement composite formulations against *S. aureus*, *S. epidermidis*, and *E. coli* through persistence Zone-of-Inhibition studies. Antibiotic release studies were also completed. To evaluate the capacity of the PMMA cement to be refilled at later time points (after initial implantation), antibiotic refilling studies were completed to simulate diffusion of drug through tissue. Finally, the porosity and mechanical properties of different formulations were evaluated to determine if they met the standards to be used for weight-bearing applications.

## EXPERIMENTAL SECTION

### Materials

β-cyclodextrin lightly cross-linked with epichlorohydrin (β-CD) prepolymer was purchased from CycloLab (Budapest, Hungary), and then further crosslinked to make insoluble polymers as described below. Ethylene glycol diglycidyl ether was purchased from Polysciences Inc. (Warrington, PA). Rifampicin (RMP) was purchased from Research Products International (Mt. Prospect, IL). Gentamicin and tobramycin were purchased from Fisher Scientific (Pittsburgh, PA). Stryker™ Simplex^®^ HV (high viscosity) radiopaque bone cement produced by Stryker Orthopaedics (Mahwah, NJ) was purchased from eSutures. Green fluorescent protein (GFP)-labeled *Staphylococcus aureus* (*S. aureus*) stock culture was kindly provided by Dr. Edward Greenfield (Case Western Reserve University, OH). *Escherichia coli* (Migula) Castellani and Chalmers (ATCC^®^ 25922) and *Staphylococcus epidermidis* (Winslow and Winslow) Evans (ATCC^®^ 12228) stock cultures were purchased from American Type Culture Collection (ATCC, Manassas, VA). Poly(tetrafluoroethylene) (PTFE) stock for cylindrical specimen molds was purchased from McMaster-Carr (Aurora, OH). All other reagents and solvents were purchased from Fisher Scientific in the highest grade available.

### Synthesis of insoluble polymerized cyclodextrin (β-CD) microparticles

Insoluble polymerized cyclodextrin microparticles were synthesized according to a previously published methodology ^17^. Briefly, 1.6 mL of ethylene glycol diglycidyl ether was added into a solution of 1 gram of β-CD (prepolymer) dissolved in 4 mL of 0.2 M potassium hydroxide. The solution was vortexed and added into a mixture of light mineral oil (50 mL) and surfactant mixture (24% Tween 85: 76% Span 85; 750 μL total volume), heated at 60°C, and stirred for four hours. Microparticles were repeatedly washed and centrifuged in gradually increasing hydrophilic solvents (i.e. oil, hexanes, acetone, MilliQ water), frozen, lyophilized, and stored in a desiccator. The size of the microparticles was previously characterized as approximately 250 μm diameter ^26^.

### Microparticle drug filling

20 mg samples of β-CD microparticles were placed in 1.5 mL tubes. 1 mL of RMP dissolved in dimethylformamide (DMF) (25 mg/mL) was added to each tube and samples were rotated for three days on a rotisserie shaker at room temperature. Drug filled microparticles were subsequently centrifuged, washed and centrifuged three times with MilliQ water, frozen, and lyophilized for 3 days. Dried drug-filled microparticles were stored in a desiccator prior to further use.

### Synthesis of PMMA spherical beads and cylinders

Small spherical PMMA beads were synthesized by combining a 2 gram sample of Stryker™ Simplex^®^ HV radiopaque surgical grade bone cement powder and 100 mg of either gentamicin or tobramycin powder (5 wt/wt% drug/cement) (comparable to concentrations used clinically) and either 100 mg (5 wt/wt% β-CD/cement) or 200 mg (10 wt/wt% β-CD/cement) of β-CD microparticles and mixed until homogeneous. The powder mixture was reacted with 1 mL of methyl methacrylate monomer for each batch of PMMA cement (in accordance with manufacturer instructions) to form a soft dough. PMMA dough with free drug and β-CD microparticles (either filled with RMP or non-drug filled (empty)) was then formed into 6 mm diameter beads and dried at room temperature ^17^.

Cylindrical PMMA samples, used in mechanical and micro-computed tomography porosity studies, were synthesized according to a previously published protocol ^17^ Briefly, a custom made two-part PTFE mold was used to fabricate cylinders 6 mm in diameter and 12 mm in height, in accordance with ASTM F451-16 standards ^27^.

### Persistence Zone-of-Inhibition (ZOI) study

A Persistence ZOI study was primarily utilized to determine the overall duration (i.e. persistence) of antimicrobial activity possible from combinatorial antibiotic PMMA composites. Persistence ZOI testing was carried out according to a previously established protocol ^17, 28–29^. Briefly, PMMA spherical beads (with different combinations of antibiotics and amounts of β-CD microparticles) were placed on the center of a Trypticase soy agar (*S. aureus* and *S. epidermidis*) or Luria-Bertani (LB) broth agar (*E. coli*) 100 mm diameter Petri dish with 70 μL of the respective bacteria culture ^17, 28–29^. Each condition was performed in triplicate. Agar plates were incubated in separate incubators at 37°C overnight and the resultant zone of bacterial clearance surrounding each drug filled PMMA spherical bead was measured (from the edge of the bead to the edge of the bacterial clearance) using calipers and averaged. PMMA samples were transferred to a freshly seeded agar Petri dish and this measurement process was continued until the zones of clearance were no longer visible.

### Quantification and activity ZOI of drug release aliquots

A drug release study was carried out in an effort to determine the amount of drug is released over time by the combination PMMA composite and to evaluate the corresponding antimicrobial effect of the drug released. Drug activity was measured in release aliquots using a modified ZOI assay. Antibiotic release from the PMMA spherical beads (containing gentamicin or tobramycin with or without RMP-filled β-CD microparticles) was carried out by placing individual beads into 1.5 mL tubes with 1 mL phosphate buffered saline (PBS). Samples were agitated at 37°C and at set time points (i.e. 2 hours, 4 hours, 1 day, 2 days, etc.) the entire release solution (1 mL) was removed from the tube and replaced with 1 mL of fresh PBS to simulate infinite sink conditions.

The actual amount of gentamicin or tobramycin drug was quantified using a ninhydrin reaction and UV absorbance spectroscopy according to a previously established protocol ^17^ Briefly, 120 μL of ninhydrin solution (2 mg/mL in PBS) was added to each release sample (400 μL). Solutions were heated at 95°C for 15 minutes (tobramycin) or 25 minutes (gentamicin), cooled, and absorbance was measured at 400 nm using a Biotek™ 96-well plate reader (H1: Winooski, VT) ^17^. Standard curves were used for the determination of concentrations in release solutions. Release of RMP was not quantified as previous work suggested that the amount of RMP released under these conditions was negligible ^17^. This study was primarily conducted to evaluate the effects of gentamicin or tobramycin release due to the addition of CD microparticles which may impact the porosity of the system.

The antimicrobial activity of each release aliquot was simultaneously evaluated according to a previous method, also using ZOI ^17^. Briefly, 6 mm diameter filter paper disks were placed in each 1 mL drug release aliquot. The activity ZOI study was then carried out using 40µL of either *S. aureus*, *S. epidermidis*, or *E. coli* ^17,30^. Bacteria were seeded on 60 mm diameter agar plates and a drug soaked filter paper disc was placed on each plate and incubated overnight at 37°C. The zone of bacterial clearance around the filter paper disc was measured for that sample, and then drug activity compared to previously determined drug amount.

### Micro-computed tomography (micro-CT) scans and porosity quantification

Micro-CT scans were collected to evaluate the porosities of PMMA samples. Micro-CT scans were collected in accordance with a previously published protocol ^17^. Briefly, cylindrical PMMA specimen were placed individually in polypropylene tubes prior to scanning. Scans were conducted using a Siemens Inveon PET-CT scanner (Siemens Medical Solutions, Malvern, PA) controlled by Siemens Inveon Acquisition Workplace software on a PC. Each sample was scanned using identical scanning and reconstruction settings as a past protocol ^17^ Data was exported in DICOM format from Siemens Inveon Research Workplace software. Cement pore segmentation/thresholding, thresholding of solid fraction, and model export were completed in 3D Slicer software (BWH and 3D Slicer contributors, version 4.8) ^31^. An upper threshold of −200 was used for segmentation of pores and a lower threshold of 0 was used for thresholding the solid fraction. Netfabb Standard 2018 (Autodesk, Inc.) was used to calculate volumetric measurements of models built in 3D Slicer. Pore volume fraction was calculated by dividing the total volume segmented by the volume of pores.

Micro-CT scans were collected for a total of 6 PMMA groups: PMMA with only tobramycin or gentamicin, PMMA with either tobramycin or gentamicin and 10wt% empty β-CD microparticles, and PMMA with either tobramycin or gentamicin and 10wt% RMP-filled β-CD microparticles.

### Compression testing

Compression testing was performed to evaluate the mechanical properties of the combination PMMA composite. The ends of PMMA cylinders were sanded square to the cylinder axis using a drill press and wet 240 grit silicon carbide sandpaper. Dimensions of individual samples were then measured (length and diameter) to the nearest 0.01 mm using digital calipers, with diameter taken at the cylinder mid-length. Samples were visually inspected after machining and only samples that contained no visible defects larger than 0.5 mm in major diameter were considered for testing. Compression testing was completed wherever possible in accordance with ASTM F451-16 ^27^, the only difference being that samples were tested approximately 48 hours after casting instead of 24 hours to allow sufficient time for machining of the cylinder ends. Cylindrical specimens were loaded in unconfined compression at a crosshead speed of 20 mm/minute and a sampling rate of 200 Hz using a mechanical testing frame (Material Testing System MTS 810, MTS Systems Corporation) equipped with a 2,000 lbf (8896 N) load cell, and load-displacement curves were analyzed for several features including ultimate strength, elastic modulus, strain to failure, and normalized work to failure. Data was first truncated to establish the onset of sample loading as the point of zero displacement. Ultimate strength, expressed in units of MPa, was determined either by taking the local maximum on the stress-strain curve before the onset of plastic deformation, or by using the 2% offset method when a local maximum was not apparent. Modulus, expressed in units of MPa, was determined from the slope of the linear portion of the stress-strain curve prior to onset of yielding using linear regression on the points between 40-60% of the determined ultimate strength. Strain to failure, expressed as a percentage of the initial sample length, was defined as the strain value at the point that the material ultimate strength was reached. Normalized work to failure, expressed in units of J/cm^3^, was determined by taking the area under the stress-strain curve from the origin up to the strain to failure.

### PMMA antibiotic-refilling in agarose phantom model

Combination PMMA composites were refilled using an agarose phantom model to test the composite’s ability to be refilled with antibiotic not initially present in the composite. A tissue-mimicking agarose phantom model was used based on a previously established protocol ^17,32^. Briefly, agarose was dissolved in PBS (0.075 wt/vol%) and heated to a boil. 5 mL hot solution was added to each well of a Costar™ 6-well cell culture plate (Corning Life Sciences, Corning, NY) and allowed to solidify. PMMA spherical beads containing free antibiotic (either gentamicin or tobramycin) with empty β-CD microparticles (either 5 or 10 wt/wt% β-CD/cement) were placed in individual wells on top of the solidified agarose. Another 5 mL hot agarose solution was added on top of each well and allowed to solidify. A small 6 mm hole was punched in the center of the top layer of the agarose. Background absorbance of agarose gels with PMMA beads were collected using a Biotek™ 96-well plate reader (19×19 point area scan; 473 nm). A small aliquot of RMP solution (6 mg/mL in methanol) was injected into each well and the plate was covered in Parafilm and incubated at 37°C with agitation. The diffusion of RMP through the agarose into the PMMA beads was monitored over 24 and 48 hours through the collection of area absorbance scans. PMMA beads were removed from the agarose and the duration of antimicrobial activity from the refilled samples was evaluated against all three bacteria species (*S. aureus*, *S. epidermidis*, and *E. coli*) in a persistence ZOI study and the total amount of RMP refilled into each bead was quantified. Studies were all completed in triplicate.

### Quantification of antibiotic refilling efficiency

The amount of RMP that was initially filled and refilled into PMMA samples was evaluated to determine the ability of combination PMMA composites to be refilled with antibiotics. Both PMMA beads containing β-CD microparticles that were initially filled with RMP and those that contained microparticles that were refilled with RMP were dissolved in 3 mL of DMSO and agitated at 37°C for 3 days. Solutions were diluted for determination of the concentration of RMP either initially filled or refilled in the β-CD microparticles in PMMA beads using absorbance spectroscopy and standard curves at 473 nm.

### Statistical analysis

Data from all studies was displayed as the mean of each condition evaluated in triplicate (persistence Zone-of-Inhibition, drug release, activity Zone-of-Inhibition, micro-CT, agarose phantom refilling) or n ≥ 3 (compression testing) with error bars depicting the standard deviation. All statistical tests were completed in Microsoft Excel 2016. Two-tailed Student’s t-tests with unequal variances were conducted for persistence ZOI study (n = 3), drug release study (n = 3), activity ZOI study (n=3), and quantification of RMP refilling (n = 3). One-tailed Student’s t-tests with unequal variances were used to analyze the compression data (n = 3 for gentamicin control, n = 4 for tobramycin control, n = 8 for 10wt% empty CD with tobramycin, n = 9 for 10wt% RMP-filled CD with tobramycin, n = 7 for 10wt% empty CD with gentamicin, n = 10 for 10wt% RMP-filled CD with gentamicin) and the porosity data obtained from micro-CT (n = 3). P-values less than 0.05 from the analyses were considered to be statistically significant.

## RESULTS AND DISCUSSION

### Persistence Zone-of-Inhibition (ZOI) study

The duration of antimicrobial activity of PMMA samples with drug combinations (RMP-filled β-CD microparticles with either gentamicin or tobramycin) and PMMA samples containing a single drug (RMP-filled β-CD microparticles only; or either free gentamicin or tobramycin only) against *S. aureus*, *S. epidermidis*, and *E. coli* were evaluated. Figure 2 depicts the results of the persistence ZOI study.

**Figure 2:**
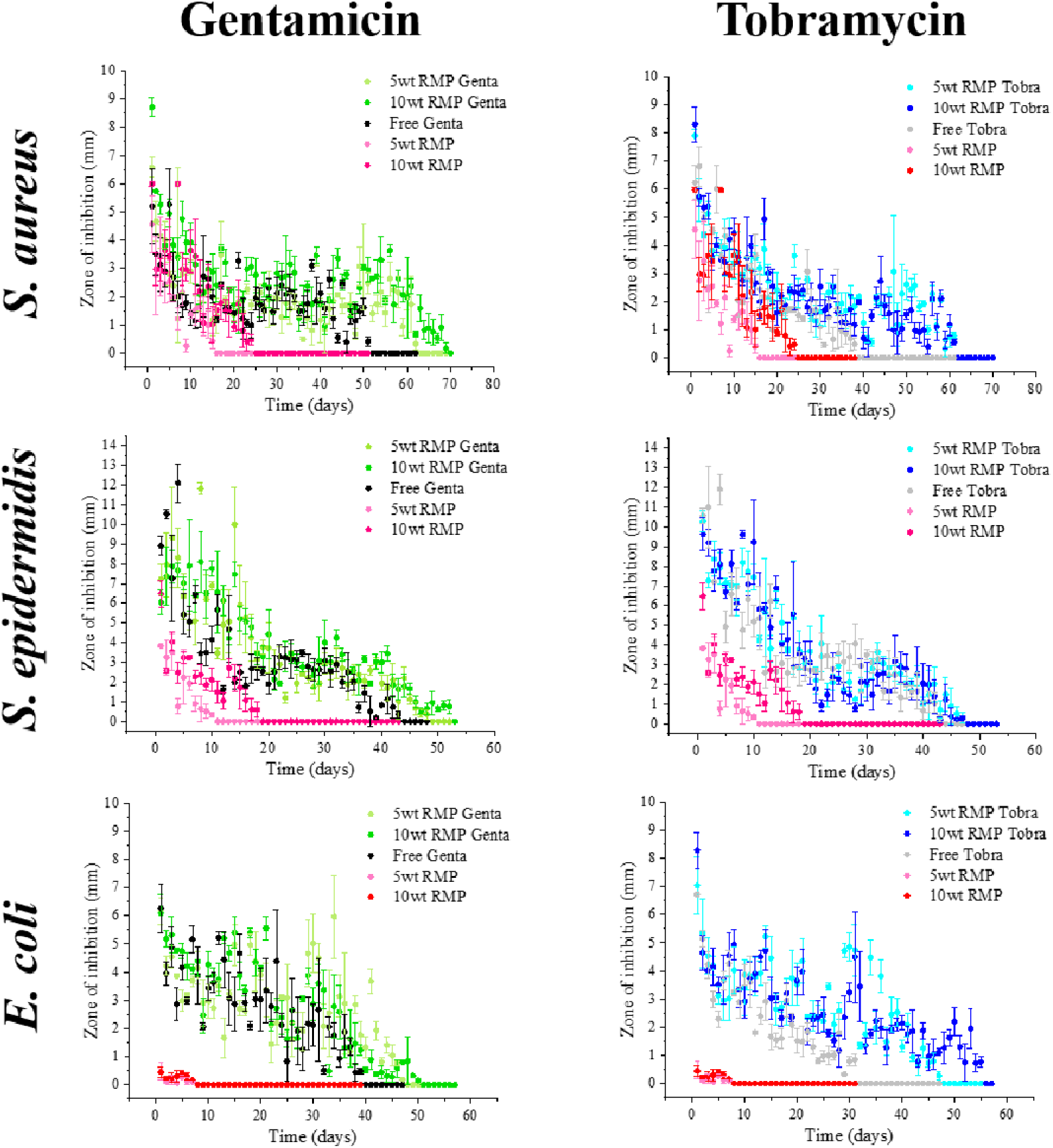
Persistence Zone-of-Inhibition studies of drug combinations (5 or 10wt% rifampin-filled CD microparticles combined with either gentamicin or tobramycin) and monotherapy drug control samples (either RMP alone; or gentamicin or tobramycin alone) against *S. aureus* (top), *S. epidermidis* (middle), and *E. coli* (bottom).

Generally, regardless of the species of bacteria, PMMA containing only RMP-filled β-CD microparticles (5 or 10wt%) exhibited the shortest duration of clearance followed by PMMA containing only gentamicin or tobramycin. Also, in all cases the longest antimicrobial effect was observed in PMMA samples containing combination antibiotics. Additionally, PMMA composite samples with two drugs typically exhibited significantly larger zones of inhibition than samples containing only RMP-filled microparticles during the early time points.

Against *S. aureus*, the longest duration of bacterial clearance was observed with PMMA containing both 10wt% RMP-filled β-CD microparticles and gentamicin lasting nearly 70 days. Specifically, during the first 10 days of the study, the drug combination samples had a significantly larger ZOI than both respective control groups (i.e. samples with only 10wt% RMP-filled β-CD microparticles or only gentamicin) (p < 0.05; except after 4, 5, 6, and 7 days where p > 0.05). Samples containing both 5wt% RMP-filled β-CD microparticles and gentamicin had only a slightly shorter duration of about 60 days. A comparable duration of activity was observed for PMMA beads containing either 5 or 10wt% RMP-filled β-CD microparticles with tobramycin (i.e. ~60 days). Specifically, over the first 10 days of the study, either 5wt% or 10wt% combination-antibiotic samples with tobramycin had a significantly larger ZOI than the respective control group (i.e. samples with only 5 or 10wt% RMP-filled β-CD microparticles) (p < 0.05; except after 3, 5, 6, and 8 days where p > 0.05). The shortest duration of clearance was observed for PMMA beads containing only RMP-filled β-CD microparticles (10wt% β-CD: 24 days; 5 wt% β-CD: 15 days). Whereas, PMMA beads containing only gentamicin or tobramycin had intermediate durations of clearance of approximately 50 and 40 days, respectively.

Against *S. epidermidis*, the longest duration of bacterial clearance was again observed with a drug combination: PMMA containing both 10wt% RMP-filled β-CD microparticles and gentamicin, lasting about 50 days. Samples containing gentamicin and 5wt% RMP-filled β-CD microparticles had a relatively comparable duration of activity as those with single drug gentamicin and 10wt% microparticles. Specifically, over the first 20 days of the study, either 5 or 10wt% drug combination samples with gentamicin had a significantly larger ZOI than the respective control group (i.e. samples with only 5 or 10wt% RMP-filled β-CD microparticles) (p < 0.05; except after 1 and 6 days for 10wt% RMP-filled microparticles where p > 0.05). Activity was also similar for samples containing both tobramycin and either 5 or 10wt% RMP-filled β-CD microparticles lasting about 50 days. Conversely, samples containing only either 5 or 10wt% RMP-filled β-CD microparticles demonstrated a dramatic decrease in the duration of activity compared to drug combination PMMA samples, specifically only about 10 and 18 days, respectively. To elaborate, over the first 20 days of the study, either 5 or 10wt% drug combination samples with tobramycin had a significantly larger ZOI than the respective control group (i.e. samples with only 5 or 10wt% RMP-filled β-CD microparticles) (p < 0.05; except after 12, 15, and 17 days where p > 0.05). As with *S. aureus*, samples containing only gentamicin or tobramycin had intermediate durations of activity around 40 days.

For *E. coli*, the longest duration of bacterial clearance was determined to be drug combination samples with 10wt% RMP-filled β-CD microparticles and tobramycin, lasing nearly 55 days. A slightly shorter duration was observed in drug combination containing samples with 10wt% RMP-filled β-CD microparticles and gentamicin (i.e. 50 days). Both samples with 5wt% RMP-filled β-CD microparticles and either free gentamicin or tobramycin demonstrated similar durations of clearance lasting nearly 50 days. In stark contrast, monotherapy samples containing only 5 or 10wt% RMP-filled β-CD microparticles only demonstrated clearance for about 7 days. Specifically, over the first 10 days of the study, drug combination containing samples with either 5 or 10wt% RMP-filled β-CD microparticles and with gentamicin or tobramycin had a significantly larger ZOI than the respective control group (i.e. monotherapy samples with only 5 or 10wt% RMP-filled β-CD microparticles) (p < 0.05). Samples with only gentamicin or tobramycin had a clearance duration of approximately 39 and 31 days, respectively.

### Quantification and activity ZOI of drug release aliquots

After the persistence Zone-of-Inhibition studies were performed, the amount and resultant activity of released drug was determined to see if it correlated with ZOI persistence. The amount of released gentamicin or tobramycin was assessed by quantifying at discrete time points for PMMA beads using a ninhydrin reaction. Simultaneously, the antimicrobial activity of each respective release aliquot (for each time point) was also evaluated using an activity ZOI study against *S. aureus*, *S. epidermidis*, and *E. coli* to ensure that the amount of drug released at each time point was capable of clearing all three species of bacteria. Daily tobramycin or gentamicin release plots and their respective antimicrobial activity are depicted in Figure 3. For each condition, there is a stacked graph where the top is the activity ZOI study and the bottom is the corresponding quantification of the daily amount of either gentamicin or tobramycin released from each sample. The quantification of drug released was normalized by dividing the mass of the drug released by the mass of the PMMA sample and was plotted in a semi-log scale. Generally, both the activity and release plots for each condition demonstrate a similar downward trend. Specifically, the activity ZOI study matches the daily amount of drug released in the sense that when a higher concentration of drug is detected in the release solution, a larger sized activity results.

**Figure 3:**
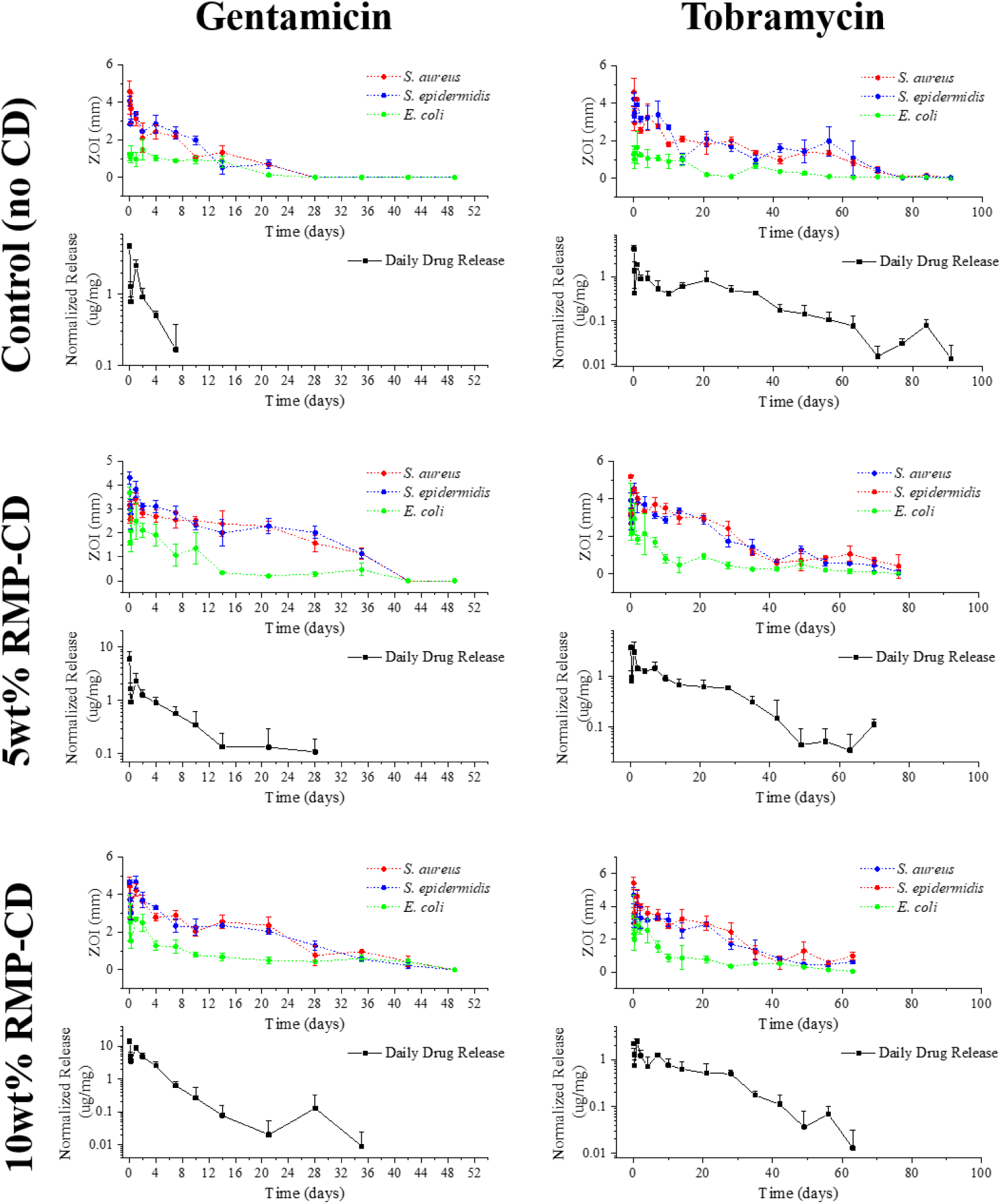
Gentamicin and tobramycin daily drug release and subsequent antimicrobial activity (based on activity Zone-of-Inhibition) of each release aliquot against *S. aureus* (red), *S. epidermidis* (blue), and *E. coli* (green) from PMMA beads containing no β-CD microparticles (top), 5wt% RMP-filled β-CD microparticles (middle), and 10wt% RMP-filled β-CD microparticles (bottom).

In general, the addition of RMP-filled β-CD microparticles to PMMA resulted in an increase in the duration of the release of gentamicin from PMMA. Specifically, samples containing only gentamicin (without any microparticles) released the drug for only 7 days. Whereas, PMMA containing either 5 or 10wt% RMP-filled β-CD microparticles in addition to gentamicin resulted in gentamicin release lasting 28 and 35 days, respectively. The amount of gentamicin released from PMMA samples containing 10wt% RMP-filled β-CD microparticles was significantly greater than the amount released from the PMMA control samples containing only gentamicin (without any microparticles) after 1, 2, and 4 days (p < 0.05). Whereas, there were no significant differences found in the amount of gentamicin released from PMMA samples containing both 5wt% RMP-filled β-CD microparticles and gentamicin to those containing only gentamicin (p > 0.05).

Correspondingly, the activity ZOI study of aliquots from PMMA containing both gentamicin and β-CD microparticles demonstrated that the aliquots had a longer duration of clearance when compared to PMMA samples with only gentamicin. More specifically, PMMA samples with only gentamicin had a clearance time of up to 28 days for all three bacteria species. Whereas, PMMA samples containing both RMP-filled β-CD microparticles and gentamicin had clearance times lasting 42 (5wt%) and 49 days (10wt%).

In contrast to PMMA samples with gentamicin, the addition of RMP-filled β-CD microparticles into PMMA did not result in an increase in the duration of the release of tobramycin from PMMA. When freely added into PMMA without β-CD microparticles, tobramycin was released over 91 days. Conversely, tobramycin was only released for 63 or 70 days when 10 or 5wt% RMP-filled β-CD microparticles were added to the PMMA. The corresponding activity ZOI study aligned with these findings. Specifically, there was a longer clearance time for PMMA samples containing only tobramycin (91 days) compared to those containing both tobramycin and RMP-filled β-CD microparticles (70 days: 5wt% β-CD; 63 days: 10wt% β-CD). No statistical significance was found between samples containing monotherapy tobramycin and drug combination samples containing either 5 or 10wt% RMP-filled β-CD microparticles with tobramycin (p > 0.05, except after 1 and 7 days in drug combination samples with 10wt% RMP-filled β-CD microparticles and after 2 and 7 days in drug combination samples with 5wt% RMP-filled β-CD microparticles).

### Micro-CT scans and porosity quantification

Micro-CT scans were collected in triplicate for a total of 6 groups of PMMA samples in an attempt to elucidate how the addition of β-CD microparticles and free drug may affect the sample porosity. Scans were analyzed to quantify the pore volume fraction, average pore volume, number of pores, and solid fraction voxel mean and standard deviation. Results are compiled in Table 1 and representative 3-dimensional renderings of the solid and pore volume fractions are depicted in Figure 4. It is important to note that the β-CD microparticles were co-registered with air (pores) during segmentation (radiodensity pure β-CD polymer = −400 HU, upper pore segmentation threshold = −200 HU; data not shown), therefore, the results that were observed regarding the porosity were partially due to the β-CD microparticles themselves.

**Table 1:** Quantification of various parameters of cylindrical PMMA samples containing either only gentamicin or tobramycin (controls), 10wt% empty (non-drug filled) β-CD microparticles with gentamicin or tobramycin, and 10wt% RMP-filled β-CD microparticles with gentamicin or tobramycin that were micro-CT scanned.^1^ Statistical significant difference of samples are relative to free tobramycin (†) and free gentamicin (‡).

| **Micro-CT Group (n = 3)** | **Pore Volume Fraction (%)** | **Average Pore Volume (nL)** | **Average # Discrete Pores** | **Solid Fraction Voxel Mean (HU)** | **Solid Fraction Voxel St Dev (HU)** |
| --- | --- | --- | --- | --- | --- |
| *Free tobra control* | 0.655 ± 0.309 | 0.531 ± 0.161 | 3362 ± 712 | 1418±57 | 488±20 |
| *Free genta control* | 0.853 ± 0.313 | 0.609 ± 0.171 | 4101 ± 569 | 1408±24 | 486±13 |
| *10wt% empty β-CD w/free tobra* | 1.797 ± 0.089† | 0.940 ± 0.120† | 5589 ± 575† | 1297±21† | 503±11 |
| *10wt% RMP-filled β-CD w/free tobra* | 1.006 ± 0.686 | 0.540 ± 0.244 | 4787 ± 1195 | 1352±22 | 489±3 |
| *10wt% empty β-CD w/free genta* | 2.381 ± 0.367‡ | 1.020 ± 0.224‡ | 6825 ± 503‡ | 1211±52‡ | 477±26 |
| *10wt% RMP-filled β-CD w/free genta* | 1.302 ± 0.187 | 0.621 ± 0.039 | 5970 ± 450‡ | 1266±38‡ | 487±13 |

<sup>1</sup> Parameters evaluated included pore volume fraction, average pore volume, number of discrete pores, solid fraction voxel mean, and solid fraction voxel standard deviation.

**Figure 4:**
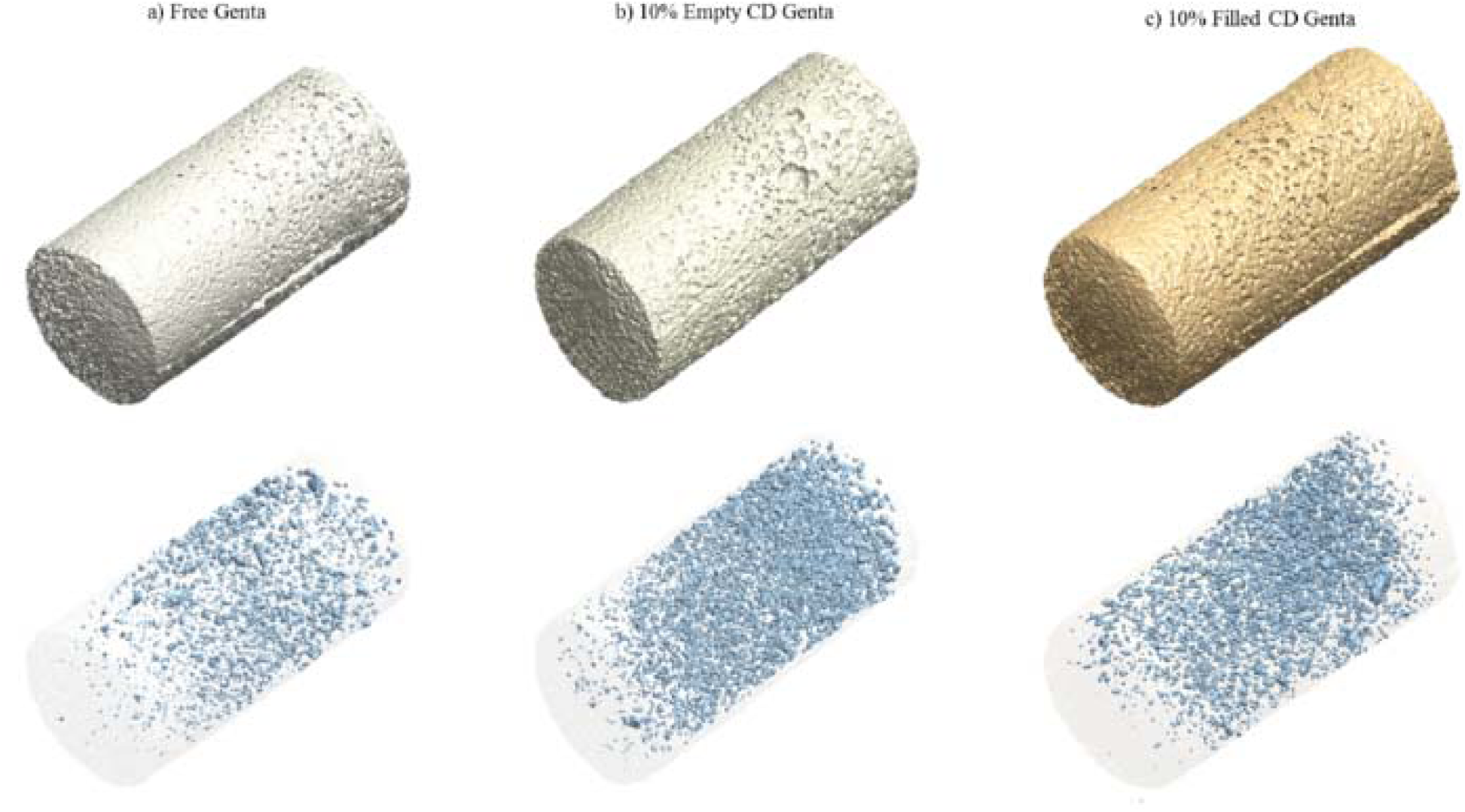
Representative 3-dimensional renderings of the solid and pore volume fractions of PMMA cylinders containing a) free gentamicin (control), b) 10wt% empty (non-drug filled) β-CD microparticles with free gentamicin, and c) 10wt% RMP-filled β-CD microparticles with free gentamicin. All were collected via micro-computed tomography scans.

Generally, the addition of β-CD microparticles increased the pore volume fraction of PMMA samples. Specifically, for PMMA containing only gentamicin, the pore volume fraction was 0.9%. Whereas, with PMMA containing both 10wt% empty β-CD microparticles and gentamicin the pore volume fraction was 2.4% and similarly for PMMA containing both 10wt% RMP-filled β-CD microparticles and gentamicin the pore volume fraction was 1.3%. A similar trend was observed in PMMA samples containing tobramycin. With tobramycin alone, the pore volume fraction was 0.7%. However, when β-CD microparticles were added it increased to 1.8% with 10wt% empty microparticles and 1.0% with 10wt% RMP-filled microparticles. Statistical analyses revealed that the differences in pore volume fractions between samples containing only gentamicin and samples containing both gentamicin and 10wt% empty β-CD microparticles as well as between samples containing only tobramycin and those containing both tobramycin and 10wt% empty β-CD microparticles were significant (p < 0.05). No significant differences were found in the pore volume fraction between the samples containing monotherapy drug (tobramycin or gentamicin) and samples containing the same corresponding drug in addition to 10wt% RMP-filled β-CD microparticles (gentamicin-filled PMMA cement: p = 0.058; tobramycin-filled PMMA cement: p = 0.241).

A similar trend was observed with average pore volume where the addition of β-CD microparticles generally increased the average pore volume. For PMMA samples containing only gentamicin, the average pore volume was 0.609 nL and significantly increased to 1.02 nL through the addition of 10wt% empty β-CD microparticles (p < 0.05). A slight, insignificant, increase to 0.621 nL was found with the addition of 10wt% RMP-filled β-CD microparticles into gentamicin-loaded PMMA samples (p > 0.05). Similarly, PMMA samples with tobramycin only had an average pore volume of 0.531 nL which increased significantly to 0.940 nL through the addition of 10wt% empty β-CD microparticles (p < 0.05). Samples containing both tobramycin and 10wt% RMP-filled β-CD microparticles had an average pore volume of 0.540 nL which was determined to be insignificant relative to samples containing only tobramycin (p > 0.05).

Addition of β-CD microparticles also generally increased the number of pores in each sample. For PMMA samples containing only tobramycin, an average of 3362 discrete pores were found in each sample. Through the addition of 10wt% empty β-CD microparticles into samples containing tobramycin, the average number of discrete pores increased significantly to 5589 (p < 0.05). A slight, insignificant, increase was also observed in the number of pores to 4787 discrete pores when 10wt% RMP-filled β-CD microparticles were added into PMMA containing tobramycin (p > 0.05). For PMMA samples containing only gentamicin, an average of 4101 discrete pores were found in each sample. Through the addition of 10wt% β-CD microparticles, the average number of pores increased significantly to 6825 discrete bodies with empty β-CD microparticles (p < 0.05) and to 5970 with RMP-filled β-CD microparticles (p < 0.05).

Solid fraction voxel mean provides a description of the average radiodensity of the solid material (i.e. excluding pores) in a given PMMA composite formulation. Our previous work has shown that the addition of RMP resulted in an increase in the solid fraction voxel mean. In general, in this study, our results suggested that incorporation of microparticles (either empty or drug-filled) into PMMA cement already containing drug reduced the average radiodensity of the solid fraction (p < 0.05), the only exception being the composite containing both 10wt% RMP-filled β-CD microparticles and tobramycin (p = 0.086). The choice of drug (tobramycin or gentamicin) when freely incorporated into non-particle-laden PMMA cement did not appear to affect the average radiodensity of the solid fraction (p = 0.399), although interestingly for particle-laden PMMA cements, use of free tobramycin was associated with a higher average radiodensity than the use of free gentamicin when the condition of particle-filling is held constant (p < 0.05). Filling of microparticles with RMP tended to increase the average radiodensity of the solid fraction (tobramycin-filled PMMA cement: p = 0.018; gentamicin-filled PMMA cement: p = 0.111).

Solid fraction voxel standard deviation provides a description of the inconsistency in the volumetric distribution of the individual solid components (especially the radiopacifier component barium sulfate) within the PMMA. Most samples demonstrated similar standard deviations around 480. This suggested that all of the PMMA composite formulations investigated possessed a similar spatial heterogeneity (p > 0.05).

### Compression testing

Cylindrical PMMA specimens of the same formulations that were CT-scanned were tested in unconfined compression to determine the effect of the presence of β-CD microparticles and free drug on the mechanical properties of PMMA. Findings are summarized in Table 2, and representative (i.e. closest to the average values for ultimate compressive strength and modulus) stress-strain curves are shown in Figure 5.

**Table 2:**
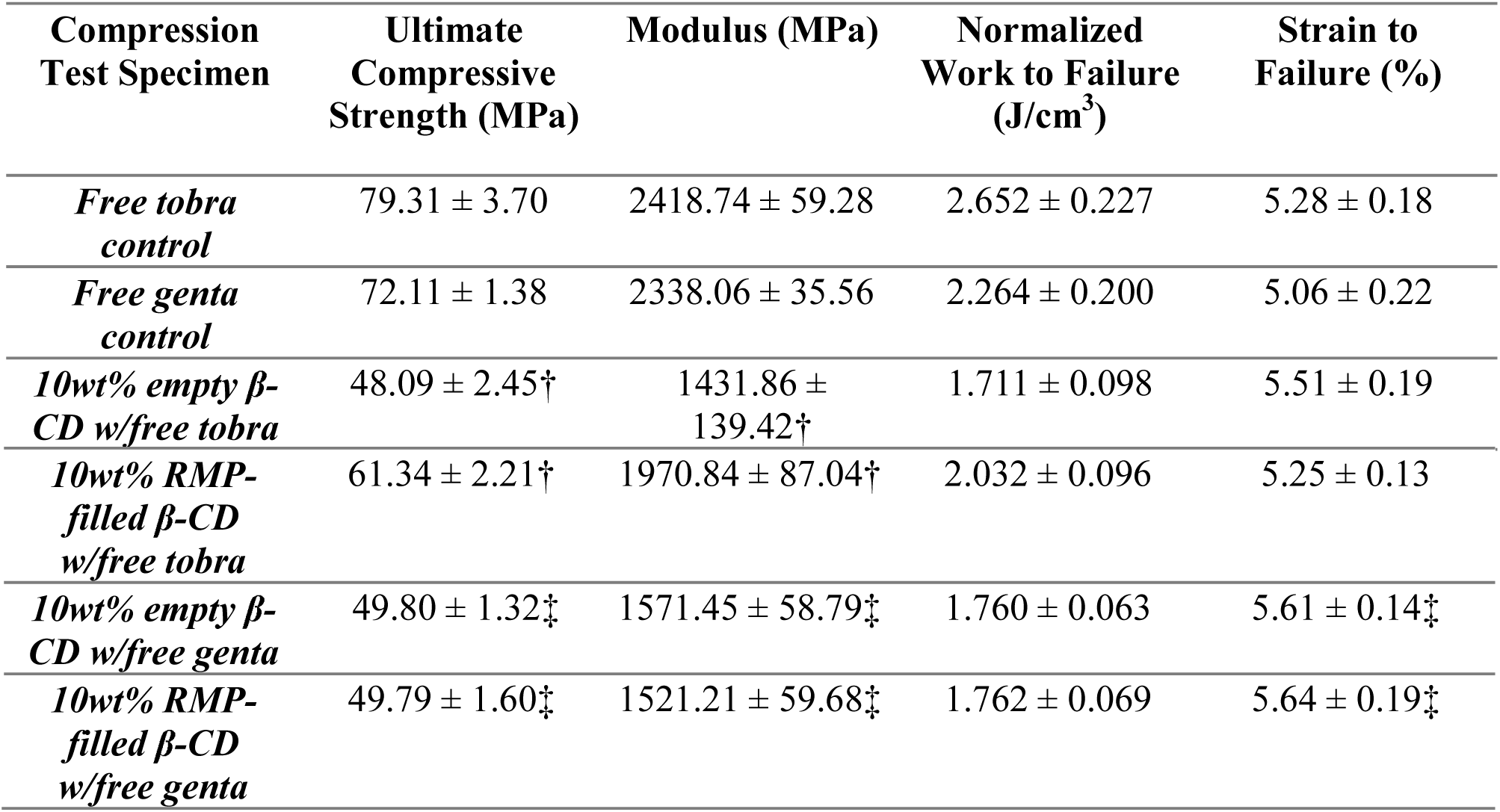
Quantification of the mechanical properties of PMMA cylindrical samples containing either a) tobramycin or gentamicin alone (controls), b) 10wt% empty (non-drug filled) β-CD microparticles with either free gentamicin or tobramycin, or c) 10wt% RMP-filled β-CD microparticles with either free gentamicin or tobramycin. All samples were evaluated via uniaxial compression.^2^ Statistically significant difference of samples relative to free tobramycin (†) and free gentamicin (‡).

**Figure 5:**
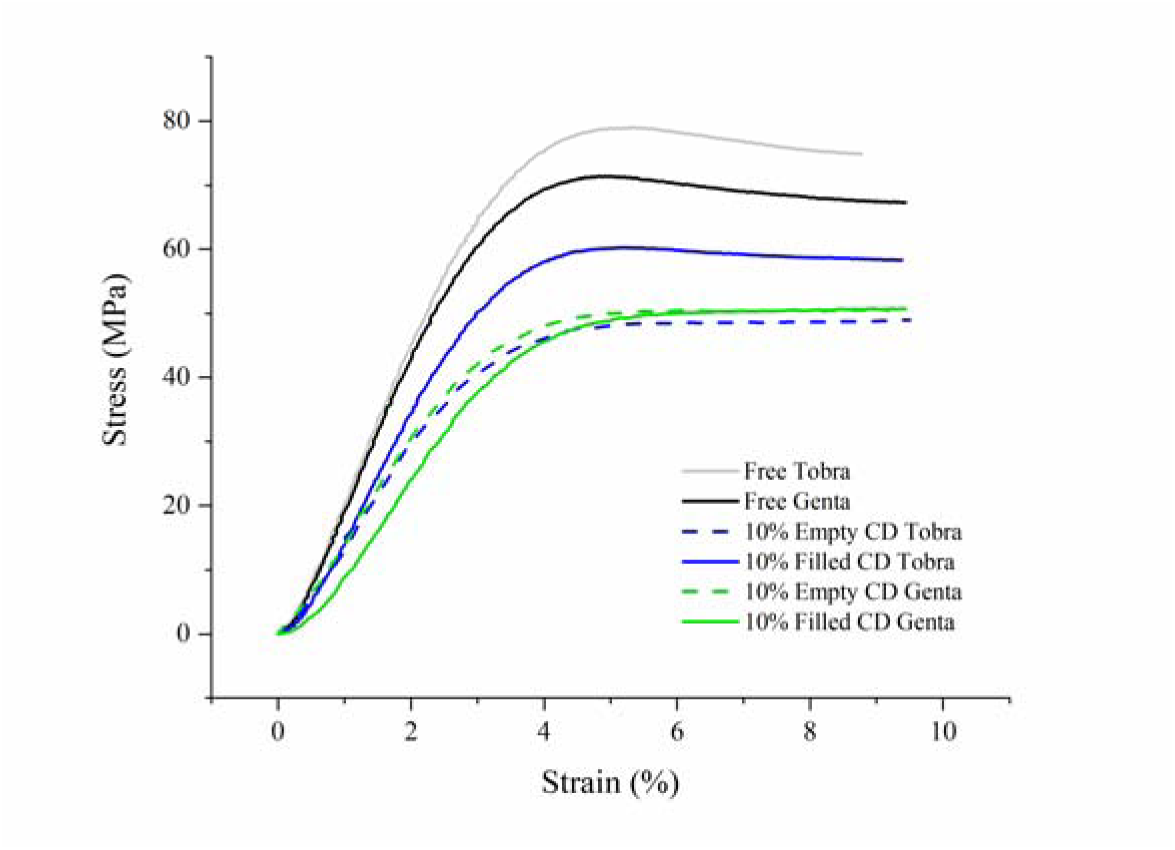
Representative stress versus strain curves from compression testing of samples.

Free gentamicin and tobramycin formulations both exceeded 70 MPa ultimate compressive strength, while all other groups were below that (~61 MPa for 10% RMP-filled CD with tobramycin, ~49 MPa for all others). For reference, plain PMMA, fabricated and evaluated in the same way but without antibiotics or additives, has an ultimate compressive strength of nearly 80 MPa (data not shown).

PMMA containing free tobramycin had a higher ultimate compressive strength (p = 0.012), modulus (p = 0.038), and normalized work to failure (p = 0.032) than PMMA containing free gentamicin, with no significant difference in strain to failure (p = 0.11). This is interesting given the structural similarity of the two drugs, but these differences may not be of major functional consequence. Addition of CD microparticles, whether RMP-filled or empty, significantly decreased compressive strength (p < 0.001), modulus (p < 0.001), and normalized work to failure (p < 0.005) of PMMA containing free tobramycin, but with no significant effect on strain to failure (p > 0.05). Likewise, addition of microparticles whether RMP-filled or empty, significantly lowered compressive strength (p < 0.001), modulus (p < 0.001), and normalized work to failure (p < 0.025), while increasing strain to failure (p < 0.02) of PMMA containing free gentamicin. Incorporating RMP into microparticles significantly increased compressive strength (p < 0.001), modulus (p < 0.001), and normalized work to failure (p < 0.001), while decreasing strain to failure (p = 0.004) of microparticle-laden PMMA containing free tobramycin whereas incorporation of RMP into CD microparticles demonstrated no significant effect on compressive strength (p = 0.493), modulus (p = 0.054), normalized work to failure (p = 0.482), or strain to failure (p = 0.343) of microparticle-laden PMMA containing free gentamicin.

### Refilling and quantification in agarose phantom model

PMMA samples containing both free tobramycin or gentamicin and empty β-CD microparticles were refilled with RMP using the tissue-mimicking agarose phantom model ^17, 32^. Figure 6 depicts images of the refilled samples after 48 hours. Table 3 displays the quantification of the amount of RMP refilled into the samples containing β-CD microparticles. Quantification was carried out on samples that were either initially empty and refilled in the agarose phantom model or that contained β-CD microparticles that were initially filled with RMP prior to being added to the PMMA. DMSO was used to dissolve away the PMMA, allowing for RMP quantification.

**Figure 6:**
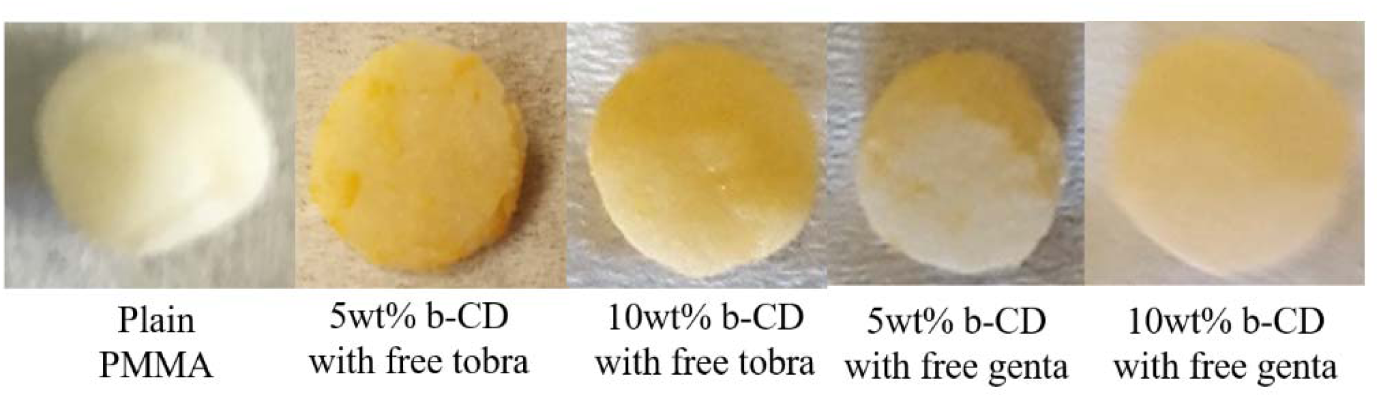
Images of refilled PMMA beads containing either no β-CD microparticles (plain) or 5 or 10wt% β-CD microparticles with either tobramycin or gentamicin after 48 hours of being refilled with RMP in the agarose phantom model.

**Table 3:**
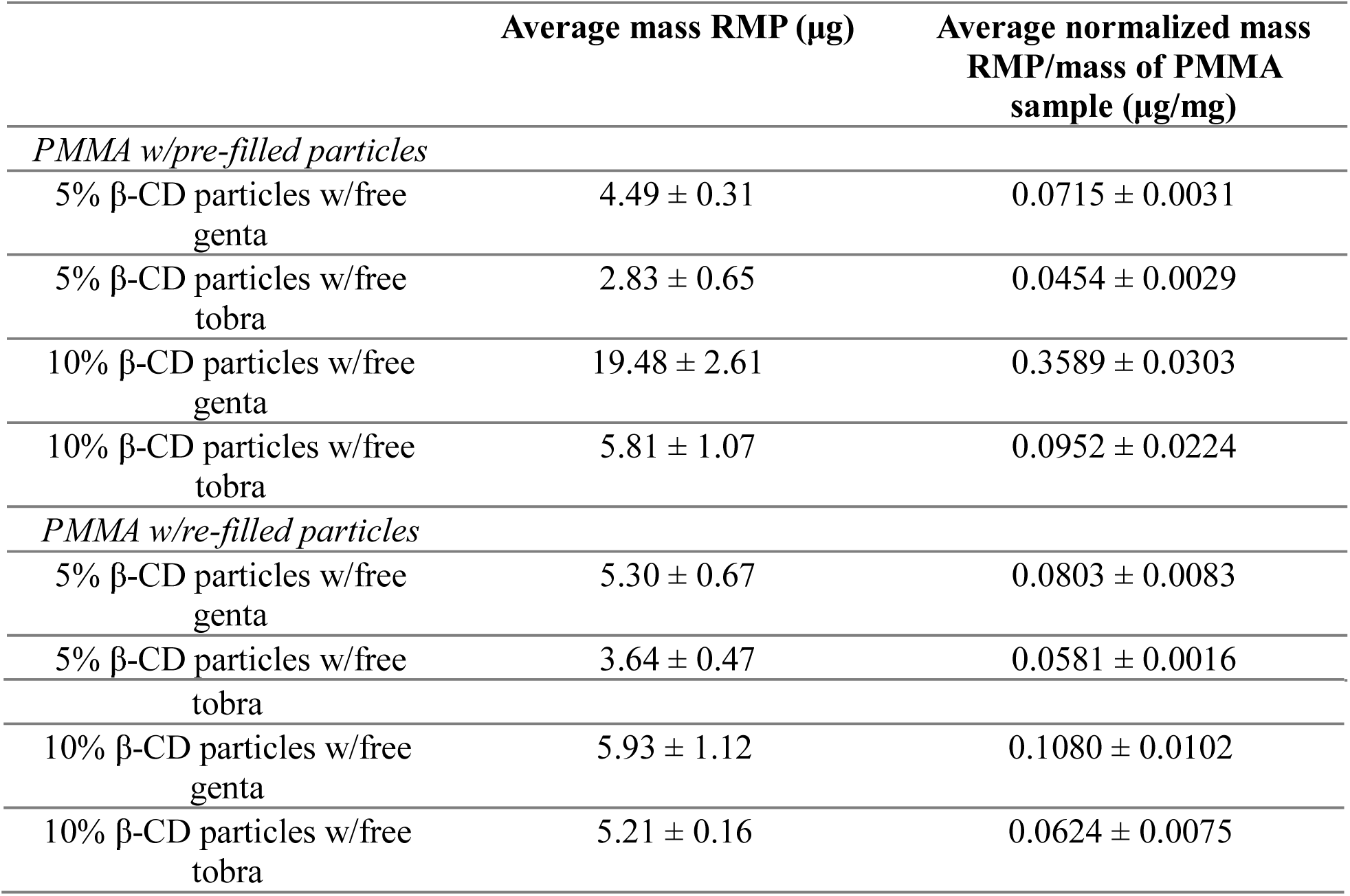
Quantification of the amount of RMP initially filled (top) and the amount of RMP re-filled through the agarose phantom model into PMMA bead samples containing either 5 or 10wt% empty (non-drug filled) β-CD microparticles with either free gentamicin or tobramycin (bottom).^3^

The experiment demonstrated the potential for refilling RMP into samples containing empty β-CD microparticles following implantation in the presence of other drugs already incorporated into PMMA (i.e. gentamicin or tobramycin).

Specifically, no statistically significant difference was found between the normalized amount of RMP loaded in pre-filled and re-filled conditions for PMMA samples containing both gentamicin and 5wt% β-CD microparticles and samples containing both tobramycin and 10wt% β-CD microparticles (p > 0.05). Similarly, there was no significant difference in the amount of normalized RMP loaded between PMMA samples containing both tobramycin and either 5 or 10wt% β-CD microparticles for both pre-filled and re-filled conditions (p > 0.05). Interestingly, samples containing both gentamicin and 10wt% pre-filled β-CD microparticles contained between three to four times more than the normalized amount of RMP relative to samples containing both tobramycin and 10wt% pre-filled β-CD microparticles.

### Persistence ZOI study (refilled PMMA beads)

Persistence ZOI studies against *S. aureus*, *S. epidermidis*, and *E. coli* were also conducted using PMMA samples that were removed from the agarose-refilling model after 48 hours (i.e. RMP refilled samples with either 5 or 10wt% β-CD microparticles and either free gentamicin or tobramycin) and the results are displayed in Figure 7.

**Figure 7:**
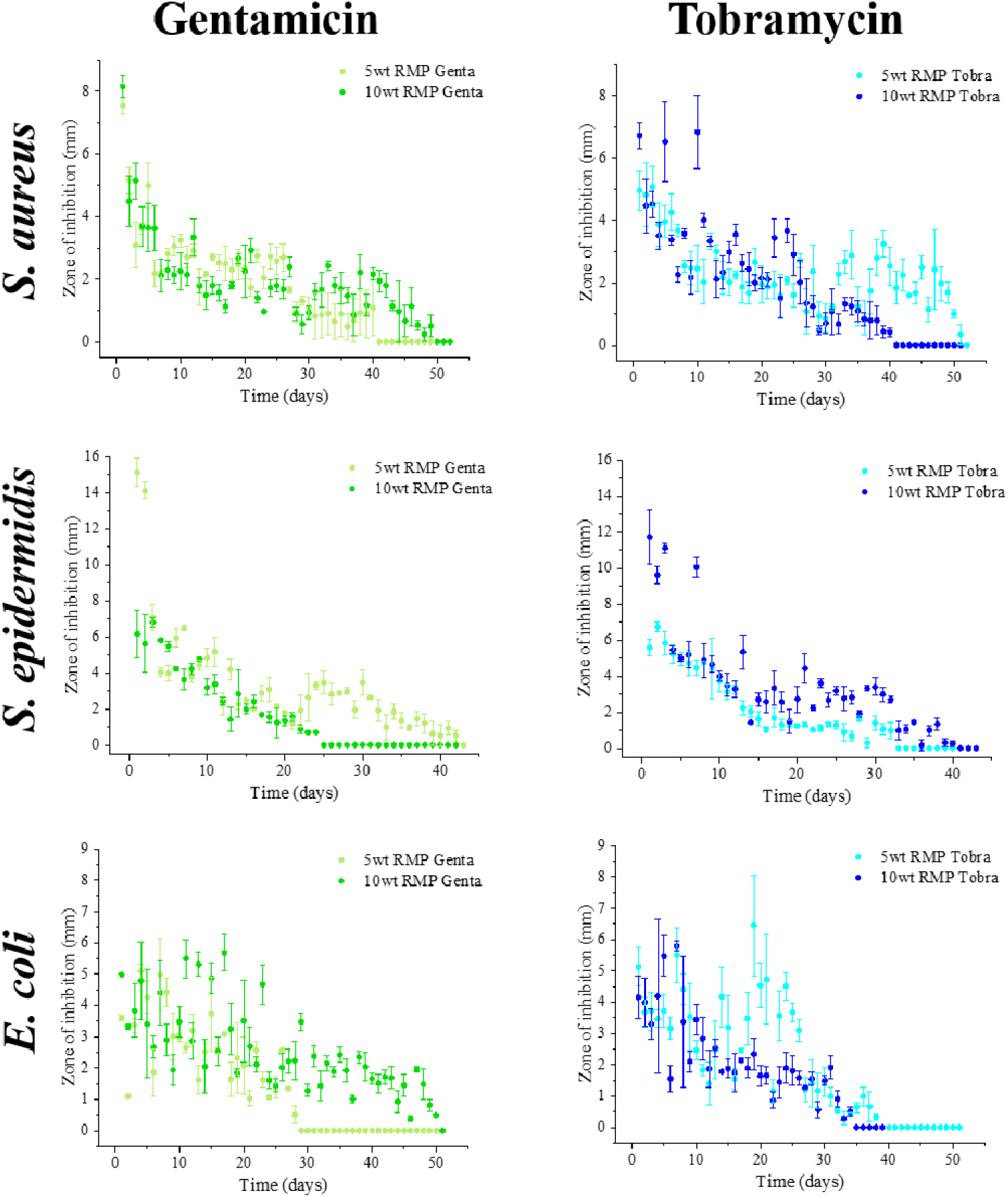
Persistence Zone-of-Inhibition studies of PMMA samples containing either 5 or 10wt% empty (non-drug filled) β-CD microparticles with either gentamicin or tobramycin that were refilled with RMP for 48 hours. Antimicrobial activity was evaluated against *S. aureus* (top), *S. epidermidis* (middle), and *E. coli* (bottom).

In general, refilled PMMA samples containing either 5 or 10wt% empty β-CD microparticle were able to allow for an additional window of antimicrobial therapy beyond what can be obtained from samples containing either gentamicin or tobramycin with 5 or 10wt% RMP pre-filled β-CD microparticles.

More specifically, against, *S. aureus*, refilled samples with gentamicin or tobramycin and β-CD microparticles had a clearance duration between 40 to 51 days. These samples also had durations of clearance between 24 and 42 days against *S. epidermidis*. Against *E. coli*, refilled samples containing either gentamicin or tobramycin with β-CD microparticles demonstrated antimicrobial activity duration ranging from 28 to 50 days.

## DISCUSSION

Different amounts of β-CD microparticles were added to clinically used gentamicin or tobramycin-laden PMMA bone cement in an effort to create a PMMA composite system to add additional antibiotic delivery potential, and more effectively treat sustained or delayed broad-spectrum infections. The premise behind our drug combination delivery system is that it allows for an enhanced antimicrobial activity by reducing the probability for antibiotic resistance to occur and by increasing the antimicrobial activity duration compared to that of a single-drug monotherapy system (i.e. for only gentamicin, tobramycin, or RMP-filled β-CD microparticles). The results supported the primary hypothesis that our drug combination composite system was able to achieve a longer antimicrobial duration against a broader range of pathogens and to be refilled with new antibiotics following simulated implantation relative to clinically-used monotherapy drug PMMA.

The increase in duration of antimicrobial activity was observed in the persistence ZOI studies against all three species of bacteria (i.e. *S. aureus*, *S. epidermidis*, and *E. coli*) when we utilized the drug combination composite PMMA system. Specifically, for *S. aureus*, there was up to a 4-fold increase observed in the duration of antimicrobial activity between a monotherapy version of PMMA and drug combination composite PMMA. For *S. epidermidis*, this comparison became a 5-fold increase, and for *E. coli* an 8-fold increase. It is possible that this improvement could be attributed to the additional mode of action against bacteria from the second antibiotic and/or the decrease in the probability for the bacteria culture to include strains that were resistant to the therapy ^9^. There may also have been a synergistic effect between the two drugs that could have allowed for an enhanced antimicrobial activity depending upon each drug’s interaction with the bacteria ^12^. Specifically, the combination of RMP and gentamicin has been shown to have an enhanced antimicrobial effect against specific staphylococcal species ^33^.

Based upon our persistence ZOI studies, the incorporation of both gentamicin and 10wt% RMP-filled β-CD microparticles into PMMA cement had the longest antimicrobial effect against gram-positive species of bacteria (i.e. *S. aureus* and *S. epidermidis*) compared to other composite formulations evaluated. Against *E. coli*, PMMA cement containing both tobramycin and 10wt% RMP-filled β-CD microparticles had the longest duration of clearance. These findings suggest that for treating or preventing poly-microbial infections from joint replacement surgeries, PMMA composite containing both gentamicin and 10wt% RMP-filled β-CD microparticles could be used for gram-positive dominant infections and PMMA composite containing both tobramycin and 10wt% RMP-filled β-CD microparticles could be used for gram-negative dominant infections.

Another observation made from the persistence ZOI studies was the relatively minor difference between the durations of antimicrobial activity between samples containing either 5 or 10wt% RMP-filled β-CD microparticles. Based upon the results from the quantification of RMP loaded into pre-filled β-CD microparticles (see Table 3), samples with 10wt% RMP-filled microparticles had between 2 to 5 times more RMP in the composite system relative to samples containing only 5wt% RMP-filled β-CD microparticles. It is possible that the small difference in duration of activity of samples may be attributed to a nonhomogeneous distribution of RMP-filled β-CD microparticles in the samples rather than the loss of RMP’s antimicrobial activity or the duration of drug release from the microparticles. The antimicrobial activity of the samples may be limited by the amount of antibiotic located near the surface of the sample that is in direct contact with the agar plate. The proximity of the antibiotic to the surface of the sample is directly related to its ability to diffuse out. This means that the RMP-filled β-CD microparticles need to be well-distributed in the sample such that the amount of drug filled microparticles located at the surface of the sample that is in direct contact with the agar plate is distinguishable between samples containing either 5 or 10wt% microparticles. Furthermore, it is possible that the contact surface may be too small to enable differentiation between samples containing either 5 or 10wt% RMP-filled β-CD microparticles. It is unlikely that the minor difference in duration of activity would be due to the loss of RMP’s antimicrobial activity. Previous studies have demonstrated that β-CD polymerized discs loaded with RMP had a duration of antimicrobial activity against *S. aureus* for more than 200 days, signifying that there was no loss of antimicrobial activity over an extended period of time ^34^.

With the majority of drug combination composite PMMA formulations we were able to achieve between 30-60 days of antibiotic release and subsequent antimicrobial activity (see Figure 3). Around 76% of total hip arthroplasty and 68% of total knee arthroplasty surgical site infections occur within 60 days after surgery. After 90 days, the risk for surgical site infection becomes lower, with the risk decreasing approximately 2-4% each month ^35^. In order to have an efficient drug delivery system that is capable of treating bacterial infections from joint replacement surgeries, the duration of antimicrobial activity of the system should reach a minimum of 60 days. Furthermore, the system should demonstrate a broad-spectrum antimicrobial activity since nearly 16% of infections from orthopedic implants are poly-microbial ^35^.

When the duration of antibiotic release from composite PMMA samples was compared to monotherapy (clinically used) PMMA formulations (i.e. only gentamicin or tobramycin) there was a 4 to 5-fold increase in the duration of the release of gentamicin from composite samples. This could be linked to the slight but insignificant increase in pore volume fractions of PMMA composite containing both gentamicin and RMP-filled β-CD microparticles observed from micro-CT scans. In contrast to the increase in duration of release of gentamicin from composite PMMA samples, a 20% decrease in the duration of the release of tobramycin was observed in PMMA samples following the addition of RMP-filled β-CD microparticles. This change in duration of release was not as striking as the change observed in samples containing only gentamicin, which may be attributed to the insignificant difference in pore volume fractions.

Studies previously completed by other groups have indicated that porosity may be a contributing factor to the total amount of drug released and the duration of sustained drug release from PMMA systems ^36–37^. This was also observed in our studies where the increase in porosity resulted in an increased duration of the release of the antibiotic. PMMA composite samples containing both gentamicin and 10wt% RMP-filled β-CD microparticles had a greater pore volume fraction and contained a significantly greater number of pores than samples containing only gentamicin (p < 0.05). The drug release studies also demonstrated that composites containing both gentamicin and β-CD microparticles had a longer duration of release compared to PMMA containing only gentamicin. Addition of 10wt% RMP-filled β-CD microparticles into samples containing tobramycin did not significantly change the pore volume fraction nor the number of pores (p > 0.05). These results were in compliance with release studies that demonstrated that the addition of 10wt% RMP-filled β-CD microparticles did not increase the duration of release for PMMA containing tobramycin. Interestingly, linear fits with R-squared values of 0.999 were observed when pore volume fraction was plotted against the duration of drug release and when the number of pores was plotted against the duration of drug release for PMMA composites containing both 10wt% RMP-filled β-CD microparticles and tobramycin or gentamicin and samples containing only tobramycin. These high R-squared values demonstrate strong negative correlations between the pore volume fraction and duration of drug release and between the number of pores in the sample and the duration of drug release. Nevertheless, it is important to note that the porosity results are partially due to the β-CD microparticles themselves as the pores and β-CD microparticles were co-registered.

Porosity has also been shown to affect the mechanical properties of PMMA cement, such as the compressive strength. Higher porosity has been associated with a lower compressive strength ^38–39^. This general trend has also been observed when the number of pores or pore volume fraction was plotted against the compressive strength of the samples. Interestingly, a strong negative linear correlation was found when the number of pores was plotted against the compressive strength (R^2^=0.872). The correlation becomes stronger when only samples containing tobramycin are considered (R^2^=0.995). In addition to the correlation found between the number of pores and compressive strength, another negative correlation was found between pore volume fraction and compressive strength. A strong linear correlation (R^2^=0.910) was observed between the two parameters when only samples that contained tobramycin were taken into consideration.

In regards to the compressive strengths, PMMA samples that contained only gentamicin were found to be weaker than samples that contained only tobramycin. This may be due to the difference in wettability of the two drugs by the liquid monomer through hydrogen bonding. Another interesting observation was that after the addition of empty β-CD microparticles, the compressive strength difference between the two was not as striking. This may be due to an affinity-based interaction between the liquid monomer and the empty β-CD microparticles. One possible explanation for the low strength of the drug combination composite formulations could be that, in comparison to the monotherapy formulations, there was a greater volume of dry solids for the same volume of methyl methacrylate liquid monomer, making the liquid volume less able to wet all of the dry solids during mixing. This ultimately could result in polymerization of shorter (lower molecular weight) PMMA chains. If so, this weakness could be corrected by increasing the volume of liquid monomer used for mixing these composite formulations, or alternatively by incorporating fewer microparticles. It is important to note, however, that preliminary studies have indicated that the PMMA cement continues to strengthen with additional time (data not shown). Additionally, it is evident that the hand-mixing procedure in this study always resulted in incorporation of some pores (at least 0.5% volume) within the PMMA, therefore vacuum mixing (as is done in clinical settings) may further elevate the strength of our composite PMMA cement materials. Further studies may need to be done to evaluate the suitability for the composite drug combination PMMA in load-bearing applications (e.g. bone cement). Nevertheless, there are applications for antibiotic-loaded PMMA bone cement (e.g. use as antibiotic beads) that do not demand the same level of structural strength as might be desired for implant fixation ^14,16–17^.

One further aspect of the superior antimicrobial activity with our composite drug combination PMMA formulation is its inherent ability to be refilled with antibiotics following implantation through the use of β-CD microparticles. Previous studies have reported that there is a negligible amount of antibiotic refilled into PMMA samples containing no β-CD microparticles following implantation relative to what is capable of being refilled in samples containing β-CD microparticles ^17^. The agarose phantom refilling studies conducted demonstrated that the composite drug combination formulations were able to be refilled with a comparable normalized amount of antibiotic to what was initially found in pre-filled samples. The subsequent persistence ZOI study suggested that these refilled RMP composite samples also retained their antimicrobial effect and that additional windows of antimicrobial activity were achieved after implantation. Furthermore, the refilling studies indicated the potential for the PMMA drug combination composite system to be implanted in the patient with empty β-CD microparticles where the patient can be provided with a local bolus of antibiotic into the tissue, near the implant as necessary. This would allow for a patient-specific therapy and would help to minimize the patient’s unnecessary exposure to antibiotics.

Refilling of β-CD microparticles is not limited to RMP, meaning that other drugs can be added into the system post-implantation. Allowing for a patient-specific therapy where drug combinations can be customized or optimized to treat infections depending upon the species of bacteria present locally. The composite PMMA formulation with β-CD microparticles would be able to target antibiotic-resistant strains of bacteria by refilling the microparticles with antibiotics which the bacteria have not been exposed to and may decrease the need for revision surgeries due to infections from joint replacement surgeries.

## CONCLUSIONS

In this study, we developed a combination-antibiotic PMMA composite system to achieve a longer and more effective antimicrobial activity against a broader range of bacteria compared to clinically-used monotherapy PMMA formulations. We observed between a 4- to 8-fold increase in the duration of antimicrobial activity against a variety of bacteria (i.e. *S. aureus*, *S. epidermidis*, and *E. coli*) when 10wt% RMP-filled β-CD microparticles were added into antibiotic-laden PMMA. In addition to the extended duration of antimicrobial activity, our composite formulation also allowed for the refilling of new antibiotics into the system’s empty β-CD microparticles, meaning that antimicrobial activity has the potential to be prolonged and that patient-specific therapy can be achieved as new suitable antibiotics are added into the system. Finally, we have examined the compressive strength and porosity of our composite formulations and found a very small, but significant decrease in the mechanical properties and increase in porosity upon addition of β-CD microparticles. The resultant mechanical strength of the tested formulations of our combination-antibiotic composite PMMA system suggest that our system may be most suitable for non-structural applications such as a temporary spacer in two-part arthroplasty revision procedures as well as antibiotic beads and that further studies may be required to determine its use in load-bearing applications such as bone cement.

## AUTHOR INFORMATION

### Author Contributions

The manuscript was written through contributions of all authors. All authors have given approval to the final version of the manuscript.

## ACKNOWLEDGMENT

The authors acknowledge financial support through National Science Foundation (NSF) Graduate Research Fellowship Program Grant No. CON501692 (E. L. C.), Center for Stem Cell and Regenerative Medicine Undergraduate Student Summer Program (ENGAGE) (C-y. L.), Support of Undergraduate Research & Creative Endeavors (SOURCE) (D. W. M.), NIH NIAMS Ruth L. Kirschstein NRSA T32 AR007505 Training Program in Musculoskeletal Research (G. D. L.), and NIH R01GM121477 (H. A. vR.).

## ABBREVIATIONS

β-CD: beta-cyclodextrin
PMMA: poly(methyl methacrylate)
RMP: rifampicin
ZOI: Zone-of-Inhibition
S. aureus: Staphylococcus aureus
S. epidermidis: Staphylococcus epidermidis
E. coli: Escherichia coli

## Footnotes

1 Parameters evaluated included pore volume fraction, average pore volume, number of discrete pores, solid fraction voxel mean, and solid fraction voxel standard deviation.

2 Parameters evaluated included ultimate compressive strength, modulus, work to failure, and strain to failure.

3 Reported values included average mass of drug in each sample and the average mass of drug in each sample normalized by the mass of the PMMA sample.

